# Valproic Acid-Induced Changes of 4D Nuclear Morphology in Astrocyte Cells

**DOI:** 10.1101/2020.06.29.178202

**Authors:** Alexandr A. Kalinin, Xinhai Hou, Alex S. Ade, Gordon-Victor Fon, Walter Meixner, Gerald A. Higgins, Jonathan Z. Sexton, Xiang Wan, Ivo D. Dinov, Matthew J. O’Meara, Brian D. Athey

## Abstract

Histone deacetylase inhibitors, such as valproic acid (VPA), have important clinical therapeutic and cellular reprogramming applications. They induce chromatin re-organization that is associated with altered cellular morphology. However, there is a lack of comprehensive characterization of VPA-induced changes of nuclear size and shape. Here, we quantify 3D nuclear morphology of primary human astrocyte cells treated with VPA over time (hence, 4D). We compared volumetric and surface-based representations and identified seven features that jointly discriminate between normal and treated cells with 85% accuracy on day 7. From day 3, treated nuclei were more elongated and flattened and then continued to morphologically diverge from controls over time, becoming larger and more irregular. On day 7, most of the size and shape descriptors demonstrated significant differences between treated and untreated cells, including a 24% increase in volume and 6% reduction in extent (shape regularity) for treated nuclei. Overall, we show that 4D morphometry can capture how chromatin re-organization modulates the size and shape of the nucleus over time. These nuclear structural alterations may serve as a biomarker for histone (de-)acetylation events and provide insights into mechanisms of astrocytes-to-neurons reprogramming.

## Introduction

Multi-cellular organisms regulate their cell type and state by selectively exposing portions of their genome for transcription through the spatial and temporal organization of chromatin—dubbed the 4D Nucleome (Chen *et al*., 2015; Cremer *et al*., 2015; Higgins *et al*., 2015, 2017a). Structurally, 3D conformation of the genome involves 1.65 turns of DNA wrapped onto a histone octamer, creating nucleosomes linked as beads-on-a-string, which are then wrapped into chromatin fibers and higher-order loops, organizing into topologically associating domains (TADs). Finally, TADs collect into a diploid set of chromosome territories (Higgins *et al*., 2015, 2017a). Histones accommodate a range of post-translational modifications, which are controlled by epigenetic proteins that ultimately regulate the transcriptional state of the cell and mediate mechanical protection of genome by chromatin rigidity. (Yang and Seto, 2007; Stephens *et al*., 2019). Chemicals that target these proteins can be used to modulate chromatin states and the concomitant cell and nuclear morphology changes observed in human diseases (Marchion *et al*., 2005; Stephens *et al*., 2019). For example, valproic acid (VPA) is a histone deacetylase inhibitor used clinically to treat epilepsy, bipolar disorders, social phobias, and neuropathic pain (Göttlicher *et al*., 2001; Ganai *et al*., 2015). Mechanistically, VPA shifts the balance towards greater histone acetylation, DNA exposure, and chromatin decondensation; activating transcriptional programs that regulate cellular processes related to cancer (Göttlicher *et al*., 2001; Eckschlager *et al*., 2017), traumatic brain injury (Higgins *et al*., 2017b), ischemia (Pickell *et al*., 2020), as well as cellular reprogramming (Huangfu *et al*., 2008). An outstanding question, however, is to characterize how mesoscale nuclear structures such as TADs mediate effects of VPA and other epigenetic modulators on cellular morphology. However, limitations of the emerging technologies that can capture these nano- (histone marks) and meso- (nuclear architecture) scale phenomena, including optical microscopy, ATACseq, and single cell RNAseq, make it difficult to simultaneously monitor macro-scale (cellular morphology) cellular state.

Recent mechanobiology studies revealed the role of chromatin as a key regulator of nuclear shape (Uhler and Shivashankar, 2018; Stephens *et al*., 2019). VPA induces increased euchromatin, which results in weakened nuclear rigidity and morphology changes that occur independently of lamins (Stephens *et al*., 2017, 2018). Here we aim to leverage this effect for assessing how the epigenetic and mesoscale nuclear state can be characterized by the overall nuclear size and shape to probe VPA’s mechanism of action. Importantly, we show that nuclear morphology can be captured in standard microscopy experiments to relate epigenetic regulators with phenotypic measurements of a cell and its organelles.

Towards this goal, we present a detailed characterization of the size and shape of astrocyte nuclei in the context of VPA treatment. We chose astrocytes because they developmentally originate from the same precursor cells as neurons and proliferate in response to brain damage (Amamoto and Arlotta, 2014), but also can be directly reprogrammed into functional neurons by sets of small molecules—often including VPA—in the lab (Cheng *et al*., 2015; Zhang *et al*., 2015; Gao *et al*., 2017; Qin *et al*., 2017). These studies, however, are not always congruent. Some reports demonstrated that VPA inhibits (Yin *et al*., 2019) or increases reprogramming efficiency (Cheng *et al*., 2015; Zhang *et al*., 2015), while others suggest that VPA alone induces astrocyte reprogramming into neurons (Cheng *et al*., 2015). The treatment protocols, concentrations, and combinations of small molecules differ between these studies, indicating that underlying mechanisms driving the trans-differentiation process are not well understood.

Existing studies that have focused on astrocyte-to-neuron reprogramming observed changes only in cellular morphology, presented in a qualitative fashion. Others quantified VPA-induced nuclear morphological changes in non-astrocyte cells, reporting time-, dose-, and cell type-dependent response. Their results ranged from no observed changes in nuclear geometry (Ganai *et al*., 2015); to altered nuclear shape (“blebbing”), but not size (Stephens *et al*., 2017, 2018); to increased nuclear size, either after 1 hour (Felisbino *et al*., 2011, 2014, 2016) or not earlier than after 7–14 days (Kortenhorst *et al*., 2009) of VPA treatment. Moreover, these approaches were limited to basic 2D surrogates of geometric measures, often reporting few features such as crosssectional nuclear area or maximum diameter. Given that chromatin is highly organized in 3D, the effects of chromatin remodeling on nuclear morphology can only be fully described in three dimensions. Supporting this assumption, there is growing practical evidence that 3D representations allow for more accurate characterizations of cell and nuclear morphology, compared to 2D measures (Choi and Choi, 2007; Meyer *et al*., 2009; Depeursinge *et al*., 2014; Kalinin *et al*., 2018a; Medyukhina *et al*., 2020). Since single-perspective 2D images depend on object’s orientation and the focal plane, they provide only a sample of its real geometry. For example, cell nuclei that differ in volume and shape can appear similarly small and circular in optically sectioned images, thus leading to lower discriminative performance of a classifier (Choi and Choi, 2007). Besides, more comprehensive morphological representation of nuclei in 3D is more informative than resolution as a determining factor for classification performance (Meyer *et al*., 2009). Our previous results also indicated that 3D size and shape descriptors outperform their 2D counterparts in the task of nuclear morphological classification (Kalinin *et al*., 2018a).

Even when the 3D object is perfectly aligned with the focal plane, only the first two out of three principal axes (major, median, minor) can be measured from its 2D representation. This reduces accuracy of shape measures computed from size features. For example, sphericity assesses the compactness of the object, i.e., it measures how closely the global object shape resembles that of a perfect sphere, which is computed via volume and surface area or approximated using principal axes (Xu *et al*., 2009). Extent (ratio of the object volume to the bounding box volume) and solidity (ratio of the object volume to the convex hull volume) of nuclear surfaces are useful measures of the amount and size of concavities (or protrusions) in an object boundary. Other shape descriptors rely on the notion of the curvature that describes how bent the curve is around each point of a surface. They allow for measuring local shape alterations that are observed on the nuclear surface and would not be exhibited in 2D projection. Mean curvature is an extrinsic measure of 3D shape that provides a balanced measure between shape morphology and curvature magnitude (Tsagkrasoulis *et al*., 2017; Kalinin *et al*., 2018b). Gaussian curvature is an intrinsic (scale-invariant) measure of curvature that depends only on distances that are measured on the surface. Shape index and curvedness are morphometric descriptors that can capture local shape features, independently or in relation to the size of an object (Koenderink and Van Doorn, 1992). Fractal dimension is the measure of the object’s boundary complexity (Metze *et al*., 2019). Together, these features allow one to measure various aspects of shape and provide a detailed quantitative characterization of 3D object morphology.

To address the limitations of previous studies, we quantified VPA-induced changes in 4D nuclear morphology of primary human astrocyte cells. Our findings show that geometric descriptors extracted from voxel and surface-modeled representations of 3D nuclear shapes enabled accurate and interpretable characterization of time-dependent morphological changes in VPA-treated astrocytes. This allowed us to distinguish between nuclear morphological profiles of treated and normal astrocytes over time with a time-average accuracy of 82%. We showed that VPA treatment induced a time-dependent increase in nuclear size and nuclear shape irregularity in astrocytes over the course of treatment.

## Results and Discussion

### Experiment and data

In order to determine how VPA-induced alterations of chromatin structure are reflected in 4D nuclear morphology, we treated human astrocyte cells with 1.5 mM of VPA at multiple time points (days 1, 3, and 5) and obtained volumetric images of DAPI-labeled nuclei using confocal microscopy at 3 time points (days 3, 5, and 7). This provided us with 4D images (3D+time) in following conditions: normal human astrocytes (NHA) and cells treated with VPA (VPA) (**Figure 1A**). First, we used deconvolution to correct for background noise and spherical aberrations in original image volumes (**Supplementary Figure 1**). Then, we segmented individual nuclei into 3D binary voxel masks (**Figure 1B**). The number of nuclei identified after segmentation and quality control are listed in **Table 1** for each day and treatment condition. Higher numbers of untreated nuclei compared to the treated group might be due to significant inhibition of astrocyte cell growth by VPA at concentrations over 1 mM (Sasai *et al*., 2007). Details of the deconvolution, segmentation, and quality control protocols can be found in the Methods section.

**Figure 1.**
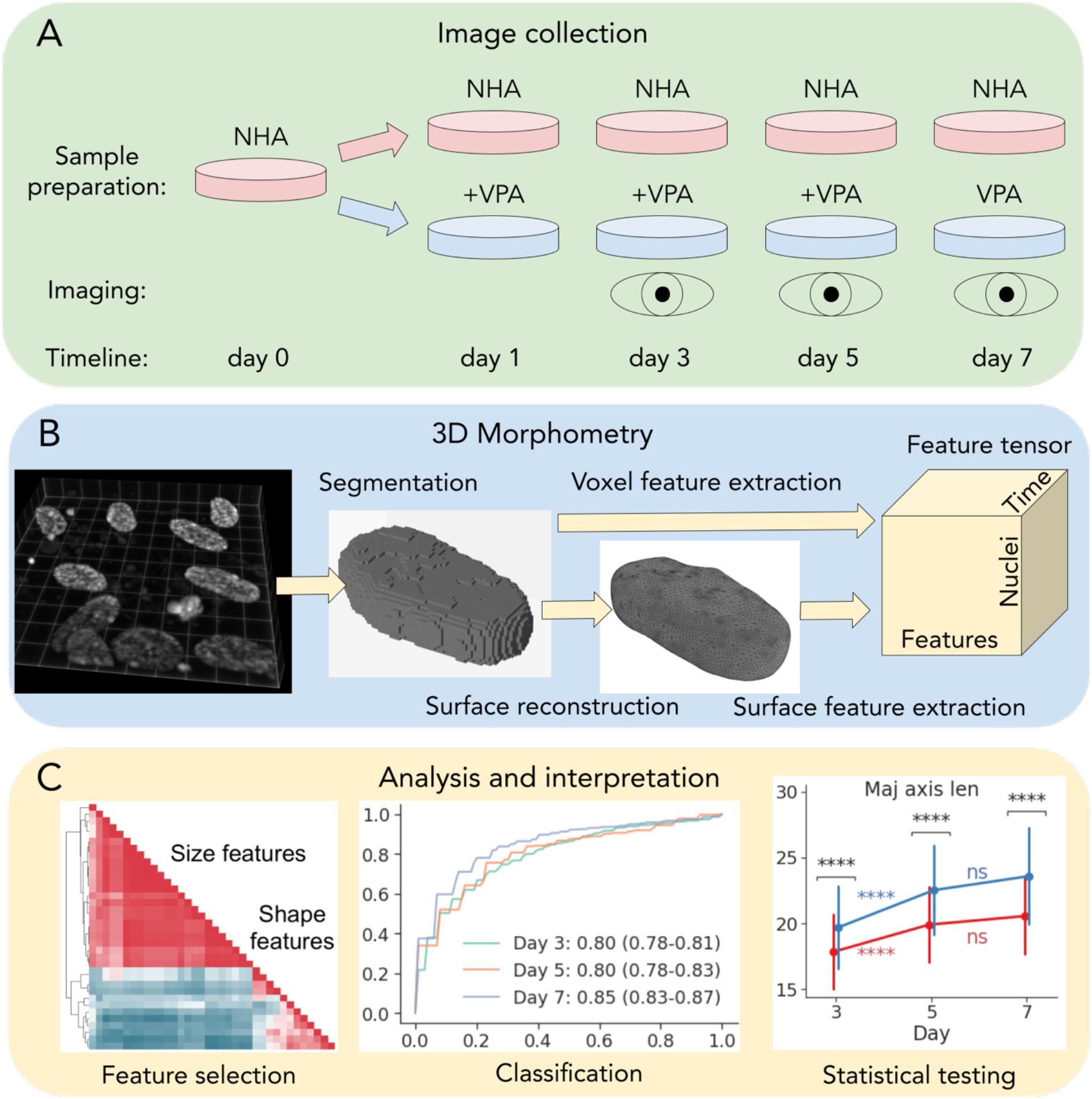
A schematic overview of the experiment, data collection and analysis. (A) sample preparation, treatment, and imaging. (B) 3D nuclear segmentation, shape modeling, and feature extraction. (C) feature selection, univariate statistical and machine learning analysis.

**Table 1.** Number of segmented astrocyte nuclear 3D binary masks per day for each treatment condition after QC (number of nuclei filtered by quality control is in parenthesis).

| Treatment | Day |  |  |
| --- | --- | --- | --- |
|  | 3 | 5 | 7 |
| NHA | 186 (-23) | 163 (-18) | 101 (-10) |
| **VPA** | 128 (-18) | 67 (-6) | 78 (-15) |
| **Total** | 314 (-41) | 230 (-24) | 179 (-35) |

A simple approach to 3D morphometry is to extract features from binary object masks represented as voxel volumes (Dufour *et al*., 2015). From each voxel binary mask (**Figure 1B**), we computed the total of 11 geometric features: volume, bounding box and convex hull volumes, extent, solidity, lengths of main axes, and inertia tensor principal components. However, voxel-based representations can be noisy and may lose fine local geometric detail or even misrepresent the object’s global topological structure. We have previously shown that nuclear surfaces obtained via shape modeling are more informative and reliable for nuclear morphometry when compared to alternatives (Kalinin *et al*., 2018b). Here, we extended that approach by extracting 16 different size and shape descriptors that can characterize morphological changes in more detail, compared to just 6 in our previous study (Kalinin *et al*., 2018b). We obtained nuclear surface representations from 3D binary masks (**Figure 1B**) and measured the same descriptors as extracted from voxels, with the addition of bounding cylinder and sphere volumes, sphericity, fractal dimension, mean and Gaussian curvatures, curvedness, and shape index. The combined feature tensor containing nuclear morphological profiles (**Figure 1B**) was used for model and feature selection, statistical and machine learning-based analysis, and interpretation (**Figure 1C**).

### Surface modeling provided compact and accurate characterization of 3D nuclear morphology

To evaluate the utility of different representations of nuclear morphology, we asked how well they facilitate morphological classification of NHA vs VPA cells, using timepoint-averaged area under the receiving operator characteristic curve (AUC) as the performance metric. Since performance of classification algorithms varies across bioinformatics problems and datasets (Olson *et al*., 2018) and not all models have feature weighting mechanism, we compared the performance across eight standard classifiers that enable feature importance estimation (**Supplementary Figure 2**). At each timepoint, we used random subsampling of the prevalent class due to high class imbalance (**Table 1**). First, since voxel and surface features have not been directly compared before in terms of nuclear morphological classification, we trained each model on sets of voxel-based (*V*) and surface-based features (*S*). As a baseline, we defined 2D voxel features extracted from 3D binary mask maximum intensity projections (*V_2d_*). To facilitate a fairer comparison between *V* and *S*, we also evaluated the subsets *V_sub_* and *S_sub_* that consisted of the 10 features that are captured by both representations (**Supplementary Figure 2**). As expected, the 2D feature set performed the worst (73% AUC), confirming that 3D measures provide more discriminative power. The performance was higher on the subset of voxel features (77% vs 74% AUC on *V_sub_* vs *S_sub_*), but not with full sets (77% AUC on *V* and on *S*). The best overall performance was achieved when using the combination of both voxel and surface-based features (78% AUC on *V*+*S*). Among tested classifiers, the support vector machine (SVM) model (Cortes and Vapnik, 1995) outperformed other classifiers averaged across all feature sets (81% AUC), so we chose to use this model going forwards.

Clustering of the complete *V*+*S* feature set revealed 2 major groups, roughly corresponding to size and shape descriptors (**Figure 2A**). To reduce redundancy and aid interpretability, we selected seven features (*S*_7_) from smaller sub-clusters that maximized SVM classification performance (82% AUC). These features included surface-based median axis length, convex hull volume, bounding sphere volume, sphericity, average mean curvature, shape index, and voxelbased solidity. *S*_7_ provided distinctive representations of nuclear morphological profiles at different time points, as shown by the t-SNE (van der Maaten and Hinton, 2008) 2D projection in **Figure 2B**. The SVM classifier demonstrated robust performance with AUCs of 80%, 80%, and 85% on days 3, 5, and 7, respectively (**Figure 2C**), with classification errors that were only slightly shifted towards false negatives at each day, indicating the effectiveness of the prevalent class subsampling (**Figure 2D**). Relative feature ranking from the trained SVM model (**Figure 2E**) revealed that sphericity was the most important feature for day 3, followed by convex hull and bounding sphere volumes. On day 5, bounding sphere volume, sphericity, and solidity were the top-3 important descriptors. Bounding sphere volume was again the most important measure on day 7, followed by average mean curvature, median axis length, and sphericity. Our findings showed that both size and shape features were important for discrimination of treated and untreated nuclei, demonstrating that a combination of 3D descriptors aids in accurate morphological classification.

**Figure 2.**
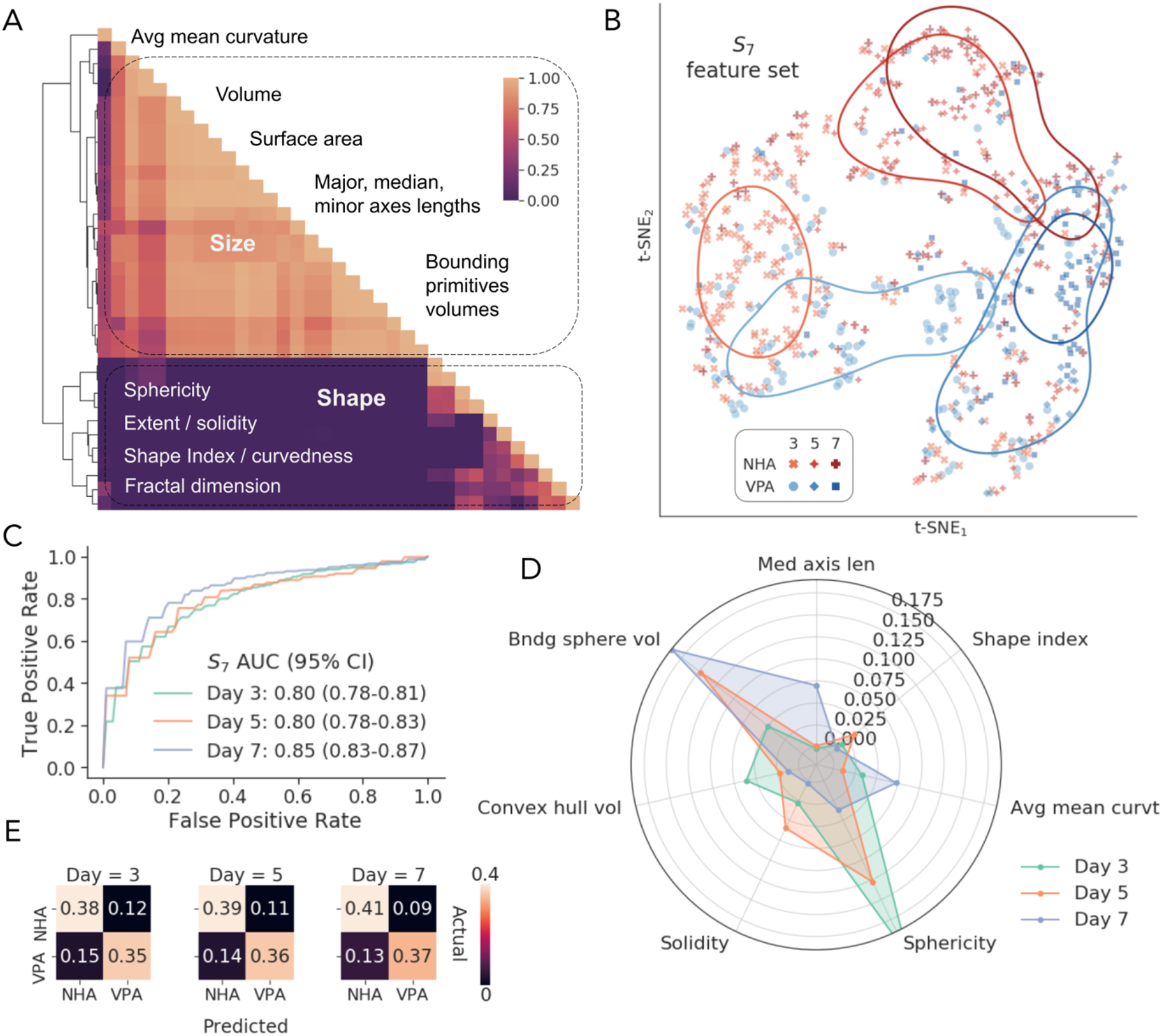
Morphological classification performance. (A) hierarchical clustering of the Pearson correlations among all voxel and surface features (*V*+*S*), showing representative size and shape descriptors. (B) 2D t-Distributed stochastic neighbor embedding (t-SNE) of the selected feature space, showing corresponding conditions (HNA or VPA) at every timepoint (day 3, 5, or 7). The lines denote clusters identified by kernel density estimation. (C) ROC curves for the SVM classifier with *S*_7_ features on days 3, 5, and 7. (D) average normalized confusion matrices for the SVM classifier on the *S*_7_ features; (E) SVM-estimated permutation importance of *S*_7_ features for distinguishing nuclear morphologies on each day.

### VPA induced increased nuclear size

We observed time-dependent alterations in astrocyte nuclear sizes as demonstrated by examples of NHA and VPA-treated reconstructed nuclear surfaces shown in **Figure 3A**, with the latter having increasingly longer major axis length and higher volume. To provide a more detailed characterization of changes in 3D nuclear size, we chose six size descriptors for further study; three of which were selected as a part of *S*_7_ (median axis length, convex hull, and bounding sphere volumes), while major and minor axes lengths and nuclear volume were chosen manually for interpretability (**Figure 3B**). Measuring the three principal axes also allowed for the generation of inferences about the global shape of nuclei, as described in the next section. We reported each relative difference as the percentage change from the mean of the control group (NHA), along with a *p*-value obtained using two-sided Mann-Whitney *U* test with Holm–Šidák multiple testing correction, and AUC that in the case of Mann-Whitney *U* test is equivalent to the common language effect size statistic (Mason and Graham, 2002).

**Figure 3.**
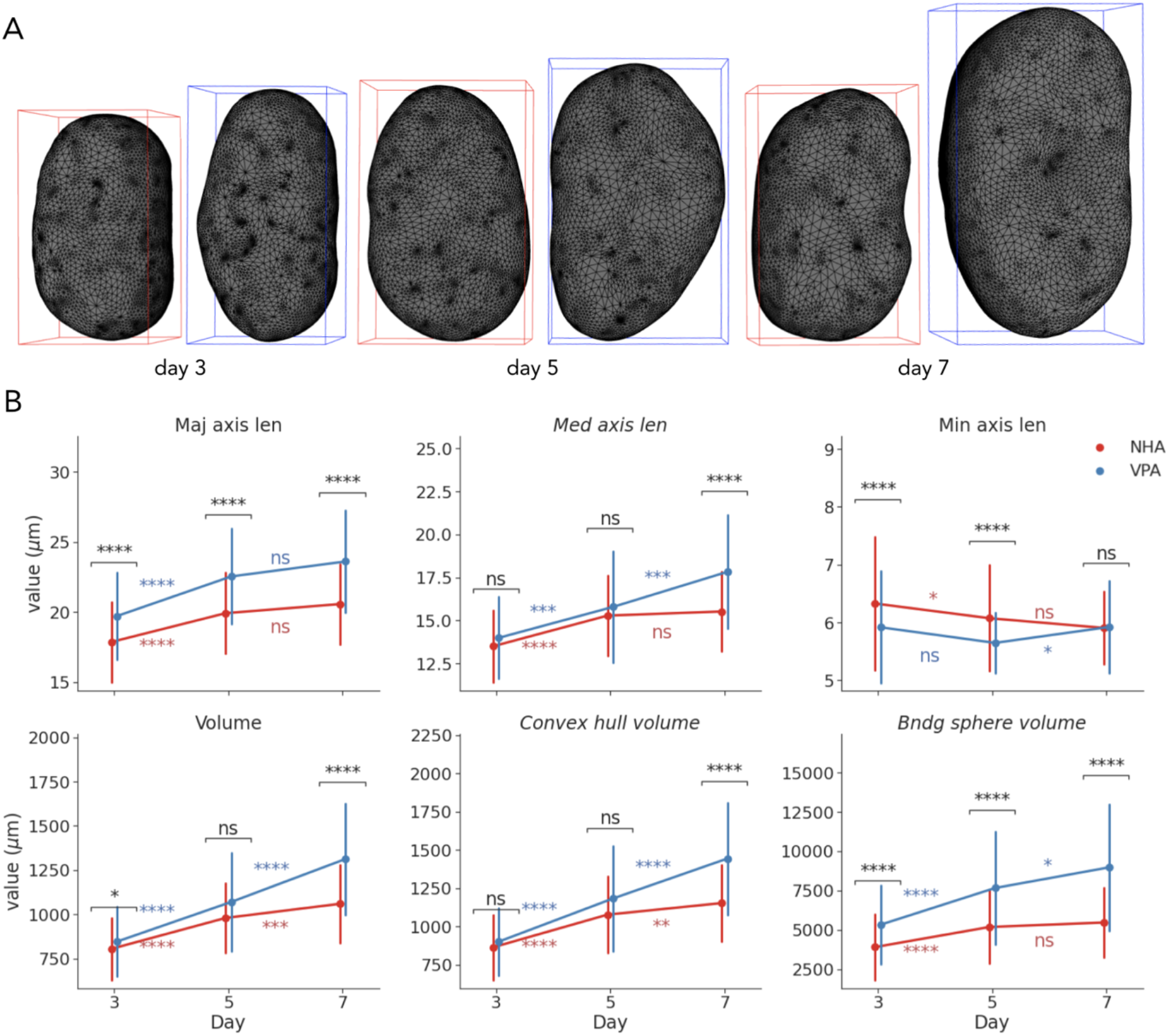
Visualization and univariate statistical analysis of size changes under VPA treatment. (A) Reconstructed surfaces of representative NHA and VPA nuclei on days 3, 5, and 7. (B) Timedependent changes in morphometric measures of nuclear sizes (points show mean; error bars show SD; \**p*<0.05, \*\**p*<0.01, \*\*\**p*<0.001, \*\*\*\**p*<0.0001).

By day 3, the average major axis length of VPA treated nuclei compared to controls had increased by 10% (*p*<0.0001, 70% AUC), the average minor axes length had decreased by 7% (*p*<0.0001, 67% AUC), while there was no significant difference in median axis lengths (**Figure 3B**). Despite more subtle changes in nuclear (+5%, *p*<0.05, 58% AUC) and convex hull volumes (+4%, *p*>0.05, 57% AUC), the volume of the bounding sphere was 36% larger (*p*<0.0001, 74% AUC) in the VPA group. Between days 3 and 5, measures of major and median axes lengths indicated intra-group changes for both NHA (+12%, *p*<0.0001 for major; +13%, *p*<0.0001 for median) and VPA nuclei (+14%, *p*<0.0001 for major; +13%, *p*<0.001 for median), along with slight shortening of minor axes. Nuclear, convex hull and bounding sphere volumes also increased in both groups, correspondingly. As the result, treated nuclei differed from controls on day 5 by having 13% longer major axis (*p*<0.0001, 73% AUC) and 4% shorter minor axis (*p*<0.0001, 67% AUC), along with 48% larger bounding sphere volumes (*p*<0.0001, 74% AUC).

Between days 5 and 7, there were no significant changes in axes lengths in the control group, while median axis length in treated group increased by 13% (*p*<0.001) and minor increased by 5% (*p*<0.05) (**Figure 3B**). By day 7, VPA-treated nuclei had 15% larger major axes (*p*<0.0001, 75% AUC) and 15% larger median axes (*p*<0.0001, 70% AUC), while minor axis length was comparable to that of untreated nuclei. Nuclear volume and convex hull volume increased in both groups from day 5 to 7, but the change in VPA-treated nuclei was more dramatic (+23%, *p*<0.0001 for volume; +22%, *p*<0.0001 for convex hull) than in the NHA group (+8%, *p*<0.001 for volume; +7%, *p*<0.01 for convex hull). As a result, the treated nuclei had 24% larger volume (*p*<0.0001, 77% AUC), 25% larger convex hull (*p*<0.0001, 76%), and 64% larger bounding sphere volume (*p*<0.0001, 80% AUC), when compared to controls on day 7.

Our findings highlight a transition point between two phases of morphological changes. Before day 5, nuclear sizes in both VPA and NHA groups significantly increased in all reported measures, except for the minor axis length. VPA-treated nuclei demonstrated more prominent elongation and flattening, along with the slightly more rapid increase of the volume, compared to controls. NHA nuclear volumes increased less between days 5 and 7, while measures of VPA-treated nuclear size continued to increase, mostly due to longer median axes, demonstrating biggest differences by the last day, as illustrated by visualizations in **Figure 3A**. Finally, all individual feature AUCs were lower than those of the SVM model at each timepoint (**Figure 2C**), which highlighted the ability of a combination of 3D linear and volumetric features to capture size alterations that are difficult to ascertain when using solely individual measures.

### VPA induced nuclear shape irregularity

Consistent with VPA decondensing the chromatin to reduce nuclear rigidity (Stephens *et al*., 2018), we observed more globally irregular surface shapes (**Figure 4 A and C**) and occasional blebbing (**Figure 4C**). As reported by the size measures (**Figure 3B**), VPA induced nuclear elongation and flattening that were reflected in more ellipsoidal and less spherical shape. To quantify, by day 3, VPA-treated nuclei demonstrated lower sphericity (−3%, *p*<0.0001, 72% AUC) and higher average mean curvature (+3%, *p*<0.0001, 71% AUC), compared to the NHA group (**Figure 4B**), which indicated the presence of more convex and less concave points on the surface (**Figure 4A**). Average Gaussian curvature and curvedness of VPA-treated nuclei were slightly lower (−5%, *p*<0.05, 58% AUC for Gaussian curvature; −5%, *p*<0.01, 59% AUC for curvedness) than those of controls, which also corresponded to overall less spherical objects. Between days 3 and 5, there were similar intra-group decreases in sphericity, shape index, and solidity that corresponded to increase in size of both treated and untreated nuclei at the same timepoint. More subtle alterations also included decrease in average Gaussian curvature in both NHA (−1%, *p*<0.05) and VPA (−5%, *p*<0.05) groups, along with the decrease of fractal dimension of the untreated nuclei (−0.2%, *p*<0.01) (**Figure 4B**). Differences between shape descriptors extracted from two conditions on day 5 were similar to those from day 3. While the difference in solidity and shape index were not identified as ‘statistically significant’ at the chosen *p*-value cut offs, they contributed to discriminative ability of the SVM classifier on both days 3 and 5 (**Figure 2D**), indicating their importance.

**Figure 4.**
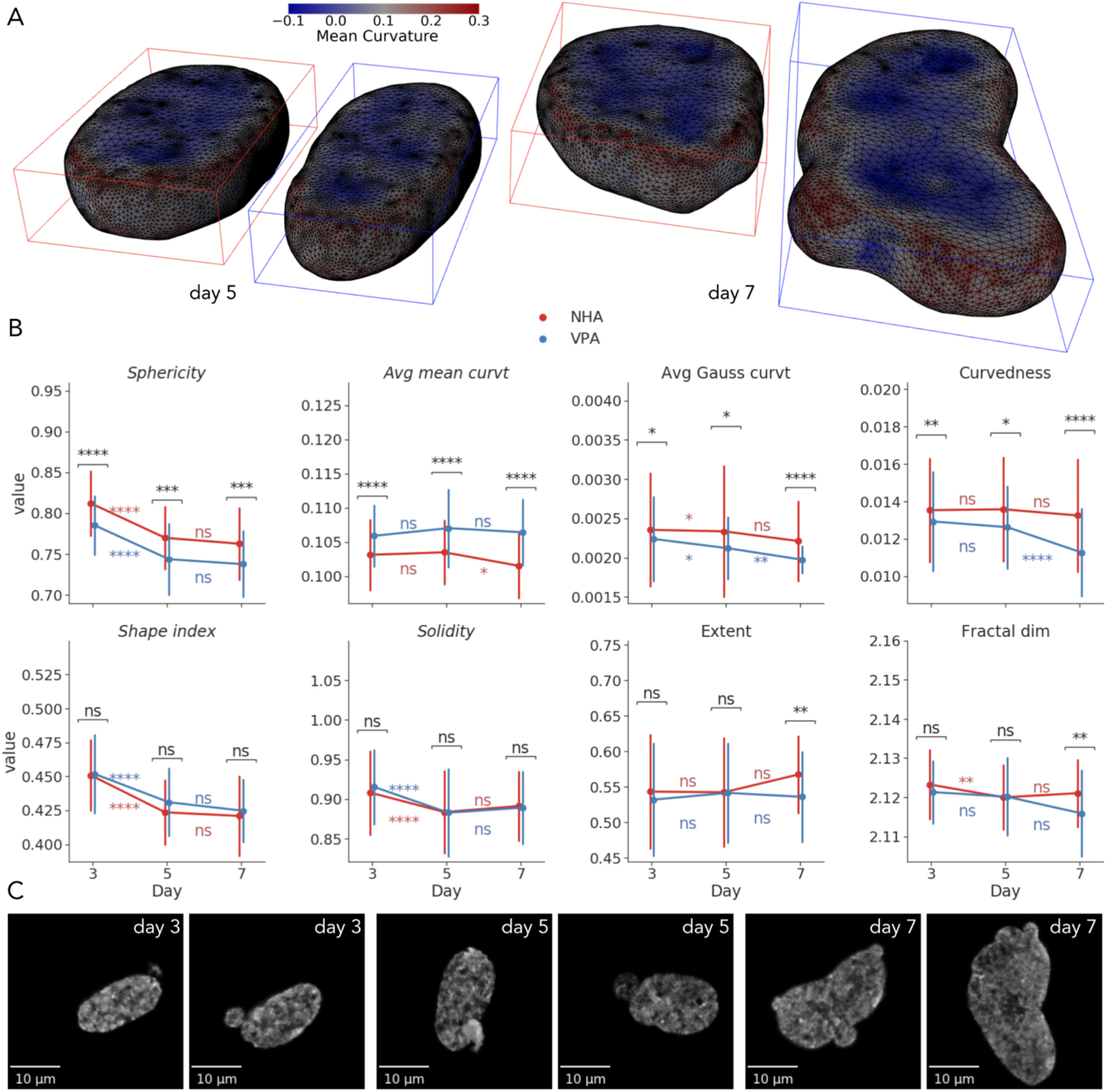
Visualization and univariate statistical analysis of shape changes under VPA treatment. (A) Reconstructed surfaces of a representative NHA and VPA nuclei on days 5 and 7, annotated with per-vertex mean curvature. (B) Time-dependent changes in morphometric measures of nuclear shapes (points show mean; error bars show SD; \**p*<0.05, \*\**p*<0.01, \*\*\**p*<0.001, \*\*\*\**p*<0.0001). (C) *XY* maximum intensity projections of VPA-treated nuclei with irregular shapes and blebbing.

By day 7, NHA nuclei only demonstrated a 2% decrease in average mean curvature (*p*<0.05) from the previous timepoint across all shape features. The VPA group exhibited an 11% decrease in curvedness (*p*<0.0001) and a 7% decrease in average Gaussian curvature (*p*<0.01) (**Figure 4B**). As on days 3 and 5, treated nuclei were still less spherical and had higher average mean curvature on day 7 compared to controls. However, decreases in curvedness (−15%, *p*<0.0001, 74% AUC) and average Gaussian curvature (−10%, *p*<0.0001, 71% AUC) were also more prominent at the last timepoint. Moreover, VPA-treated nuclei had 6% lower extent (*p*<0.01, 64% AUC), and 0.2% lower fractal dimension (*p*<0.01, 66% AUC), which indicated overall higher global shape irregularity and less intensively folded and wrinkled nuclear surfaces, respectively.

Overall, we showed that VPA robustly induced more elongated and less spherical nuclear shapes with higher mean curvature from day 3. Towards day 7, these nuclei demonstrated more prominent deformations characterized, for example, by a kidney-like form and blebbing (**Figure 4C**). At the same time, the decrease in curvedness and fractal dimension reflected nuclear surfaces with lower local border complexity that has also been suggested to be indicative of chromatin decondensation, reduced tumorigenesis, and neuroprotection (Metze *et al*., 2019). Combined with the size features, these 3D shape descriptors enable more accurate characterization of time-dependent morphological changes than their 2D counterparts or any single measure used individually.

Together, our findings described the dynamics of VPA-treated nuclear morphological profiles characterized by the increased sizes and progressively more irregular, complex shapes that can be attributed to altered histone modifications and chromatin decondensation. These observations represent a first step to studying time-dependent morphological effects of chromatin reorganization in the astrocyte-to-neuron reprogramming process and relating them to underlying molecular mechanisms. In future studies, this approach can be extended to label additional sub-nuclear components or organelles, such as nucleoli, chromosome territories, TADs, transcriptional condensates, and other compartments using, for example, Cell Painting assay(s) for high-content morphological profiling (Bray *et al*., 2016). While tracking the overall cellular phenotype, this extension will allow us to include many other features in the models and assess their variability, association with cellular and nuclear shape morphology, disease state, and treatment conditions. All of this will set the stage to evaluate the effects of VPA and other small molecules measured with different concentrations and temporal sampling. Correlating these phenotypical cell and nuclear profiles with data from other assays, such as Hi-C (Lieberman-Aiden *et al*., 2009), will likely provide useful insights into how altered functional properties of the genome are correlated with TAD structure, nuclear and cellular morphology, and can it be used for training machine learning models to more accurately predict treatment response in model systems and in humans (Kalinin *et al*., 2018c).

## 4 Methods

### Sample preparation and image acquisition

Primary human astrocyte cells were purchased from ScienCell (Human Astrocytes hippocampal, Catalog #1830).

#### Day 0

Replace media with 50% growth media and 50% N2 media (DMEM/F12 + 1X pen/strep, 1X N2 supplements).

#### Day 1

Control samples (NHA): for 30ml of N2 media add 36ul DMSO

VPA-treated (1.5mM VPA): for 30ml of N2 media add 450ul VPA

#### Day 3

Collect day 3 control and VPA samples:

- fix samples in 4% PFA for 10 min
- rinse 3 × 5 min in PBS
- store samples in PBS at 4deg

Control samples (NHA): for 30ml of N2 media add 187.5ul DMSO

VPA-treated (1.5mM VPA): for 30ml of N2 media add 450ul VPA

#### Day 5

Collect day 5 control and VPA samples:

- fix samples in 4% PFA for 10 min
- rinse 3 × 5 min in PBS
- store samples in PBS at 4deg

Control samples (NHA): for 30ml of N2 media add 37.5ul DMSO

VPA-treated (1.5mM VPA): for 30ml of N2 media add 450ul VPA

#### Day 7

Collect day 7 control and VPA samples:

- fix samples in 4% PFA for 10 min
- rinse 3 × 5 min in PBS
- store samples in PBS at 4deg

Cells in both collections were labeled with DAPI (4’,6-diamidino-2-phenylindole), a common stain for the nuclear DNA. 3D imaging employed a Zeiss LSM 710 laser scanning confocal microscope with a 63x PLAN/Apochromat 1.4NA DIC objective. Each original 3D volume was then re-sliced into a 1,024×1,024×*Z* lattice (*Z*={30,50}), where regional sub-volumes facilitate the alignment with the native tile size of the microscope. For every sub-volume, accompanying vendor metadata was extracted from the original data.

### Image pre-processing and segmentation

To correct for imaging artifacts, including the effect of axial smearing on volumetric measures, we deconvolved the obtained images. Theoretical 3D point spread functions for each individual image volume were modeled using the Richards & Wolf algorithm from the PSF Generator plugin for Fiji (Kirshner *et al*., 2013). We then used estimated point spread functions and imaging metadata to apply Lucy-Richardson deconvolution (10 iterations) to the original 3D image volumes using the DeconvolutionLab2 software (Sage *et al*., 2017). Deconvolution reduced axial smearing and improved segmentation and classification results (**Supplementary Figure 1**).

We performed the automatic 3D segmentation of nuclei from deconvolved images using Nuclear Segmentation algorithm from the Farsight toolkit (Al-Kofahi *et al*., 2010; Kalinin *et al*., 2018a). The main advantage of this tool is that it was created specifically to segment DAPI-stained nuclei in 2D or 3D. Unlike other learning-based segmentation algorithms, it does not require a labeled training set. It demonstrated stable results in our previous project (Kalinin *et al*., 2018a) and on these data. The algorithm implements multiple steps, which include a graph-cut algorithm to binarize the sub-volumes, a multi-scale Laplacian of Gaussian filter to convert the nuclei to blob masks, fast clustering to delineate the nuclei, and nuclear contour refinement using graph-cuts with alpha-expansions (Al-Kofahi *et al*., 2010).

After segmentation of the DAPI channel sub-volumes, data were converted to 16-bit 3D TIFF files, each segmented nucleus was represented as a binary mask, and given a unique index value. Post-segmentation processing of nuclear masks included 3D hole filling and a filtering step that removed the objects if they span the edge of a tile, are connected to other objects. This quality control protocol filtered out most of the artifacts and remaining were removed by visual inspection.

### 3D morphometry

3D binary nuclear masks were then used for voxel-based feature extraction using scikit-image Python library (van der Walt *et al*., 2014), obtaining feature set *V*. For each binary mask we measured nuclear volume, bounding box and convex hull volumes, extent, solidity, lengths of principal axes, inertia tensor principal components, and diameter of the sphere with the same volume as the nucleus. In order to compare performance of 2D vs 3D features, we also extracted 2D features from maximum intensity projections of binary masks.

First, we model boundaries of nuclear 3D masks extracted from the microscopic images as genus zero two-dimensional manifolds that are embedded as triangulated surfaces in 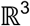 (Kalinin *et al*., 2018b). 3D surface modeling uses an iterative Laplace-Beltrami eigen-projection and a topology-preserving boundary deformation algorithm (Shi *et al*., 2010). This algorithm performs robust reconstruction of the objects’ surfaces from their segmented masks using an iterative mask filtering process. We used Mask2Mesh implementation of this algorithm from the MOCA framework (Shi *et al*., 2010) and then applied mesh simplification and subdivision to 40,000 triangles, following the shape analysis protocol in (Kalinin *et al*., 2018b). The next step included extraction of geometric characteristics of the 3D nuclear surfaces, e.g., mesh volume, surface area, curvedness, shape index, and fractal dimension, implemented in LONI Pipeline v7.0.3 (Dinov *et al*., 2009; Kalinin *et al*., 2018b). In order to provide a comprehensive morphological characterization, we expanded the list of six metrics previously reported in (Kalinin *et al*., 2018b). The additional surface features were extracted using trimesh library (Dawson-Haggerty et al., 2019) and included average mean curvature, convex hull and bounding primitive (box, oriented box, cylinder, sphere) volumes, convex hull surface area, inertia tensor eigenvalues, principal axes lengths. Surface-based extent and solidity were computed from trimesh-derived measures as the ratio of the object volume to the bounding box volume and the ratio of the object volume to the convex hull volume, correspondingly. Sphericity of the nucleus was computed as the ratio of the surface area of a sphere with the same volume as the given nucleus to the surface area of the nucleus: 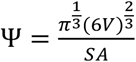, where *V* is the volume of the nucleus and *SA* is the surface area of the *SA* nucleus (Wadell, 1935). All surface descriptors were combined into the feature set *S*.

Both volumetric and surface features we merged by inner join on the per-nucleus basis, filtering out those individual cells, for which feature extraction failed or voxel volume measure exceeded the empirically estimated threshold of 150000 voxels. Nuclei were also considered outliers and removed if any feature value computed over all cells was out of the [P_5_ – 1.5(P_95_ – P_5_);P_95_ + 1.5(P_95_ – *P_5_*)] range, where *P_5_* and *P_95_* are 5^th^ and 95^th^ percentiles, correspondingly. All extracted features were aggregated into per-nucleus feature vectors, from which we constructed a feature table per each day of treatment with the labels corresponding to phenotypic conditions (NHA and VPA).

### Feature and model selection, and analysis

In order to handle multicollinear features, we performed hierarchical clustering on the Spearman rank-order correlations between all features and then used averaged SVM classification performance to select a threshold for defining feature clusters, while controlling for overfitting with 5-fold statistical cross-validation repeated from 5 different random seeds. From each cluster, we selected one feature according to the highest value for the *χ*^2^ statistic, while ignoring inertia tensor eigenvalues, as they were highly correlated with other features that are easier to directly interpret, such as volume or minimal principal axis length. This process yielded the final set *S*_7_ consisting of the following 7 descriptors: surface-based median axis length, convex hull and bounding sphere volumes, sphericity, average mean curvature, shape index, and voxel-based solidity. t-SNE embedding was generated using scikit-learn library (Pedregosa *et al*., 2011) with PCA initialization, perplexity of 13, and cosine distance as a metric. Clusters for each condition at every timepoint were identified using kernel density estimation in seaborn library (Waskom *et al*., 2020) with 0.75 threshold. It should be noted that cluster sizes and inter-cluster distances should be interpreted with care when using t-SNE (Wattenberg *et al*., 2016).

Following sets of features were used to compare volumetric and surface-based shape representations: *V_2d_*, *V*, *S*, *V_sub_*, *S_sub_*, *V*+*S*, *S*_7_, where *V_sub_* and *S_sub_* were subsets of equivalent voxel and surface feature sets correspondingly (nuclear volume, convex hull and bounding box volumes, extent, solidity, inertia eigenvalues, and major axis length). We assessed classification performance of those classifiers from with scikit-learn library (Pedregosa *et al*., 2011) that provided feature weights as an output. These included Gaussian naïve Bayes, k-nearest neighbors, logistic regression, linear support vector machine (SVM), random forest, extremely randomized trees, AdaBoost and gradient boosting machine. They were trained using default hyperparameter values and evaluated at each timepoint using the Area under the Receiving Operator Characteristic. AUCs, confusion matrices, and feature ranking were averaged from 10 repetitions of the internal statistical 5-fold stratified cross-validation with different random seeds.

We averaged the performance of each model over timepoints and over all feature sets. To compare feature sets, we averaged the AUCs over all models and days. Random subsampling of the prevalent class by the total number of per-fold training samples from an underrepresented class at each iteration of the cross-validation procedure was used to combat class imbalance. To rank features by their relative importance, we employed the permutation importance strategy that reflects the decrease in a model performance when a single feature value is randomly shuffled (Breiman, 2001). This breaks the relationship between the feature and the outcome, while being model agnostic and can be calculated many times with different permutations of the feature. We computed permutation importance on a held-out set on each cross-validation cycle, highlighting which features contribute the most to the generalization power of the trained model. Permutation feature importance was computed using the *‘coef_’* property of the trained SVM model at each timepoint.

Univariate statistical analysis of individual features was performed using SciPy package (Virtanen *et al*., 2020) with multiple testing correction using statsmodels (Seabold and Perktold, 2010). We reported each relative difference as the percentage change from the control group, along with a p-value obtained using two-sided Mann-Whitney *U* test with Holm–Šidák multiple testing correction (α=0.05), along with the difference of means (DoM), and the common language effect size statistic that in the case of Mann-Whitney U test is equivalent to the Area under the Receiving Operator Characteristic curve (AUC) (Mason and Graham, 2002).

For all analysis tasks we used Python 3.8.3 from the Anaconda distribution (Continuum Analytics and others, 2016), with *numpy* (van der Walt *et al*., 2011), pandas (McKinney, 2010), and iPython (Perez and Granger, 2007) packages. Image processing was done using scikit-image (van der Walt *et al*., 2014). Visualizations and charts were built with matplotlib (Hunter, 2007), seaborn (Waskom *et al*., 2020), trimesh (Dawson-Haggerty et al., 2019), and SOCRAT (Kalinin *et al*., 2017) libraries.

The documentation supporting the conclusions of this article together with the derived data, pipeline workflows, and underlying source code will be made publicly available online upon publication of the paper on the project webpage: SOCR 3D Cell Morphometry Project, https://socr.umich.edu/proiects/3d-cell-morphometry.

## Acknowledgements

The authors thank Ari Allyn-Feuer, Samuel Handelman, and Gilbert Omenn for thoughtful discussions and helpful suggestions. We thank colleagues at the Laboratory of Neuro Imaging (LONI), at the Keck School of Medicine, University of Southern California, for providing technical support for the LONI Pipeline environment. This work was partially supported by the National Science Foundation grants 1916425, 1734853, 1636840, 1416953, 0716055, 1023115; the National Institutes of Health grants P20 NR015331, P30 DK089503, UL1TR002240, R01CA233487, R01MH121079. Xin Rong of the University of Michigan donated NVIDIA Titan X GPU used for this research, and the NVIDIA Corporation donated TITAN Xp GPU used to execute the computationally intensive protocol.

