## Supplemental figures for "Valproic Acid-Induced Changes of 4D Nuclear Morphology in Astrocyte Cells"

Alexandr A. Kalinin *et al*.

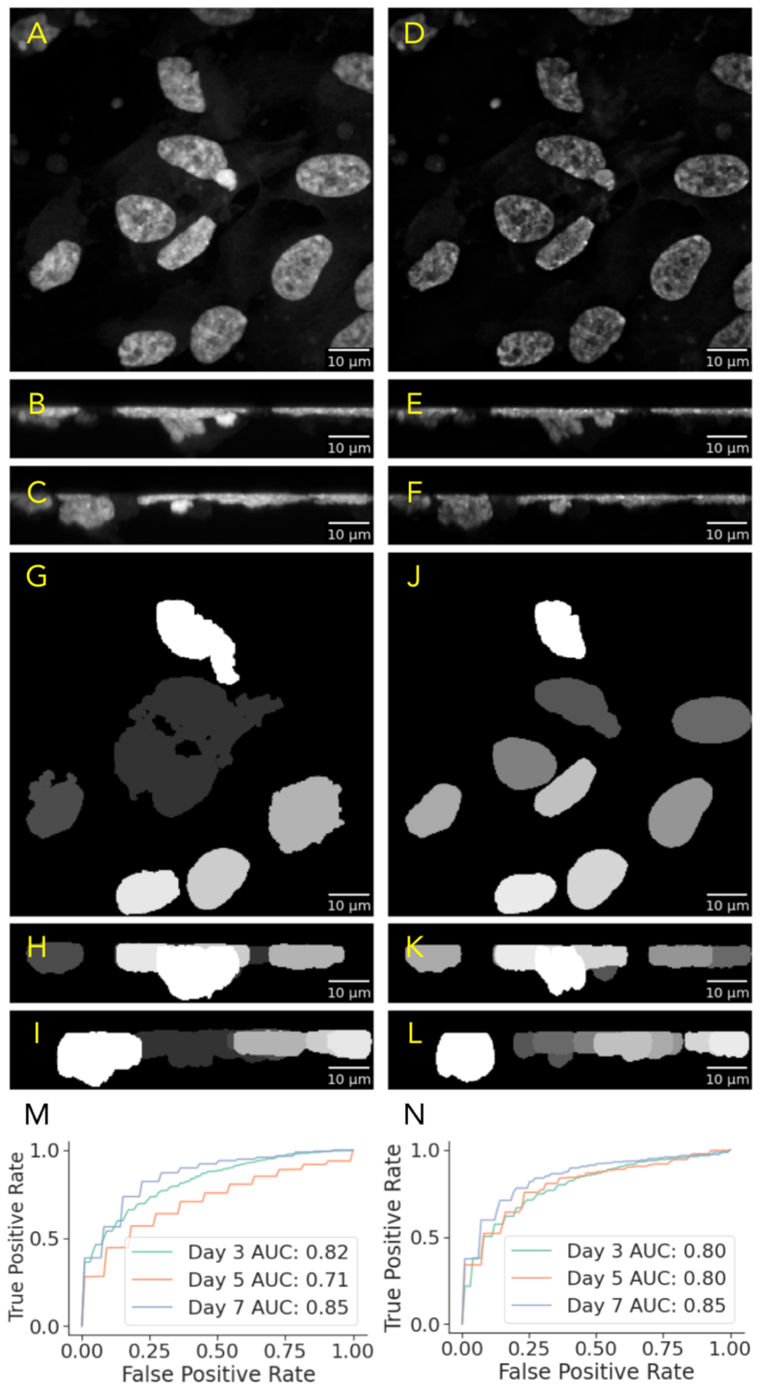


**Supplementary Figure 1**. Effects of image deconvolution on axial smearing, nuclear segmentation and classification. (A, B, C) *XY, XZ, YZ* maximum intensity projections of the original example volume. (D, E, F) *XY, XZ, YZ* maximum intensity projection of the deconvolved example volume. (G, H, I) *XY, XZ, YZ* maximum intensity projection of the segmented original example volume. (J, K, L) *XY, XZ, YZ* maximum intensity projection of the segmented deconvolved example volume.

(M) AUC curves for the best classification performance on selected features extracted from original segmentation masks (79% average AUC, Random Forest model, 11 features). (N) AUC curves for the best classification performance on selected features from deconvolved segmentation masks (82% average AUC, SVM model, 7 features).

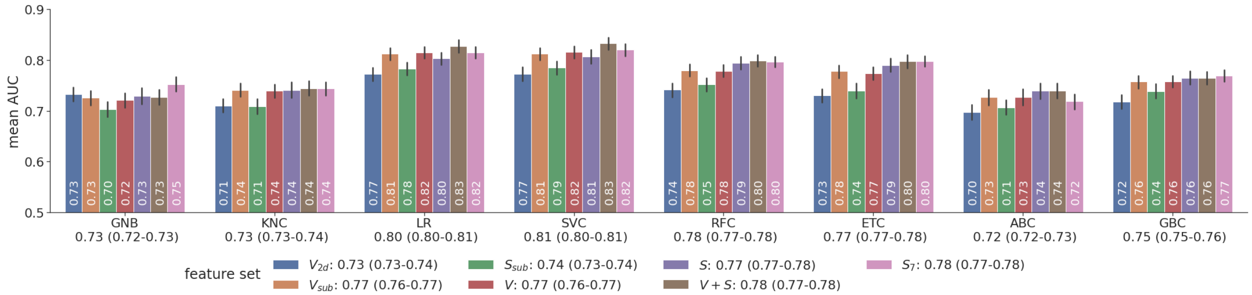


**Supplementary Figure 2**. Classifier and feature set selection. Mean AUCs (with 95% bootstrapped confidence intervals) over days 3, 5, 7 for different classifiers (GNB– Gaussian naïve Bayes; KNC–k nearest neighbors; LR–logistic regression; SVC–support vector machine; RFC–random forest; ETC–extra-randomized trees; ABC–AdaBoost; GBC–gradient boosting) and feature sets: $V_{2d}$–features of 2D projections of $V$; $V_{sub}$ and $S_{sub}$– subsets of the 10 features shared between $V$ and $S$; $V$–only voxel features; $S$–only surface features; $V$+$S$–the union of V and S features; and $S_7$–the subset of 7 features selected with hierarchical clustering.
